# Non-blocking modulation contributes to sodium channel inhibition by a covalently attached photoreactive riluzole analog

**DOI:** 10.1101/228684

**Authors:** P Lukács, M C Földi, L Valánszki, E.H Casanova, B Biri-Kovács, L Nyitray, A Málnási-Csizmadia, A Mike

## Abstract

Sodium channel inhibitor drugs decrease pathological hyperactivity in various diseases including pain syndromes, myotonia, arrhythmias, nerve injuries and epilepsies. Inhibiting pathological but not physiological activity, however, is a major challenge in drug development. Sodium channel inhibitors exert their effects by a dual action: they obstruct ion flow ("block"), and they alter the energetics of channel opening and closing ("modulation"). Ideal drugs would be modulators without blocking effect, because modulation is inherently activity-dependent, therefore selective for pathological hyperactivity. Can block and modulation be separated? It has been difficult to tell, because the effect of modulation is obscured by confromation-dependent association/dissociation of the drug. To eliminate dynamic association/dissociation, we used a photoreactive riluzole analog which could be covalently bound to the channel; and found, unexpectedly, that drug-bound channels could still conduct ions, although with modulated gating. The finding that non-blocking modulation is possible, may open a novel avenue for drug development because non-blocking modulators could be more specific in treating hyperactivity-linked diseases.

## Introduction

### Sodium channels as drug targets

Voltage-gated sodium channels (VGSC) are essential components of electrical signal propagation in excitable tissues. Dysfunction of sodium channels may cause hyperexcitability, leading to several pathologies, including different pain syndromes, certain types of epilepsy, myotonia and arrhythmia. Hyperexcitability may also ensue from modification of channel and pump functions following mechanical injury, ischemic injury or inflammation. Overexcitation is thought to be involved in several neurodegenerative and psychiatric diseases (Eijkelkamp et al., 2012; Tarnawa et al., 2007). Inhibition of sodium channels may be an effective treatment for these conditions, however, non-selective inhibition could not be beneficial because of the vital role sodium channels play in neuronal and muscle function. Isoform selective sodium channel inhibitor drugs could be a solution for this problem, but due to a highly conserved drug-binding region (Payandeh and Minor, 2015), it has been difficult to develop isoform-selective drugs (Bagal et al., 2015; England and de Groot, 2009). Fortunately, most sodium channel inhibitors exert a certain degree of functional selectivity, showing a definite preference for cells with abnormally high activity or a slightly depolarized membrane potential. In order to be able to find and develop drugs with high functional selectivity, it is essential to understand the mechanisms behind this phenomenon. Sodium channel inhibitors differ remarkably in their modes of action (Lenkey et al., 2010): which conformations they prefer, at which conformations can they access their binding site, and what are the rates of association and dissociation. We also propose in this study that they might also differ in the way inhibition is effectuated: by channel block or by modulation.

### Drug action on sodium channels: characteristics and possible underlying mechanisms

Sodium channel inhibitors can exert their effect via two major mechanisms. Channel block means physical occlusion of the pore that prevents conduction sterically or electrostatically. Modulation, on the other hand, produces inhibition by energetically stabilizing one of the channel's native non-conducting conformations. This is typically inactivated conformation, a state assumed by the channel upon prolonged depolarization (either after opening or even without opening), which is essential in preventing overexcitation, and in making signal propagation by self-regenerating sodium channel activation. Common sodium channel inhibitor drugs produce a weaker inhibition at hyperpolarized membrane potentials, which is assumed to be due to channel block, and a much stronger inhibition at depolarized membrane potentials, which is thought to be due to a higher degree of channel block and, in addition, to modulation as well. The ability to modulate by stabilizing inactivated state also implies that the drug must have higher affinity to this conformation, according to the modulated receptor hypothesis (Hille, 1977; Hondeghem and Katzung, 1977). Besides state-dependent affinity, state-dependent accessibility also contribute to the strong dependence of inhibition on membrane potential, as pointed out by the guarded receptor hypothesis (Starmer et al., 1984). The result of state-dependence is manifested in phenomena typical of sodium channel inhibitors: Besides reduced amplitude of sodium currents, the voltage dependence of availability is shifted towards hyperpolarized potentials, as measured in the widely used "steady-state inactivation" (**SSI**) protocol; and the recovery from the inactivated state is delayed, as measured in the "recovery from inactivation" (**RFI**) protocol (Fig. 1).

**Fig. 1.**
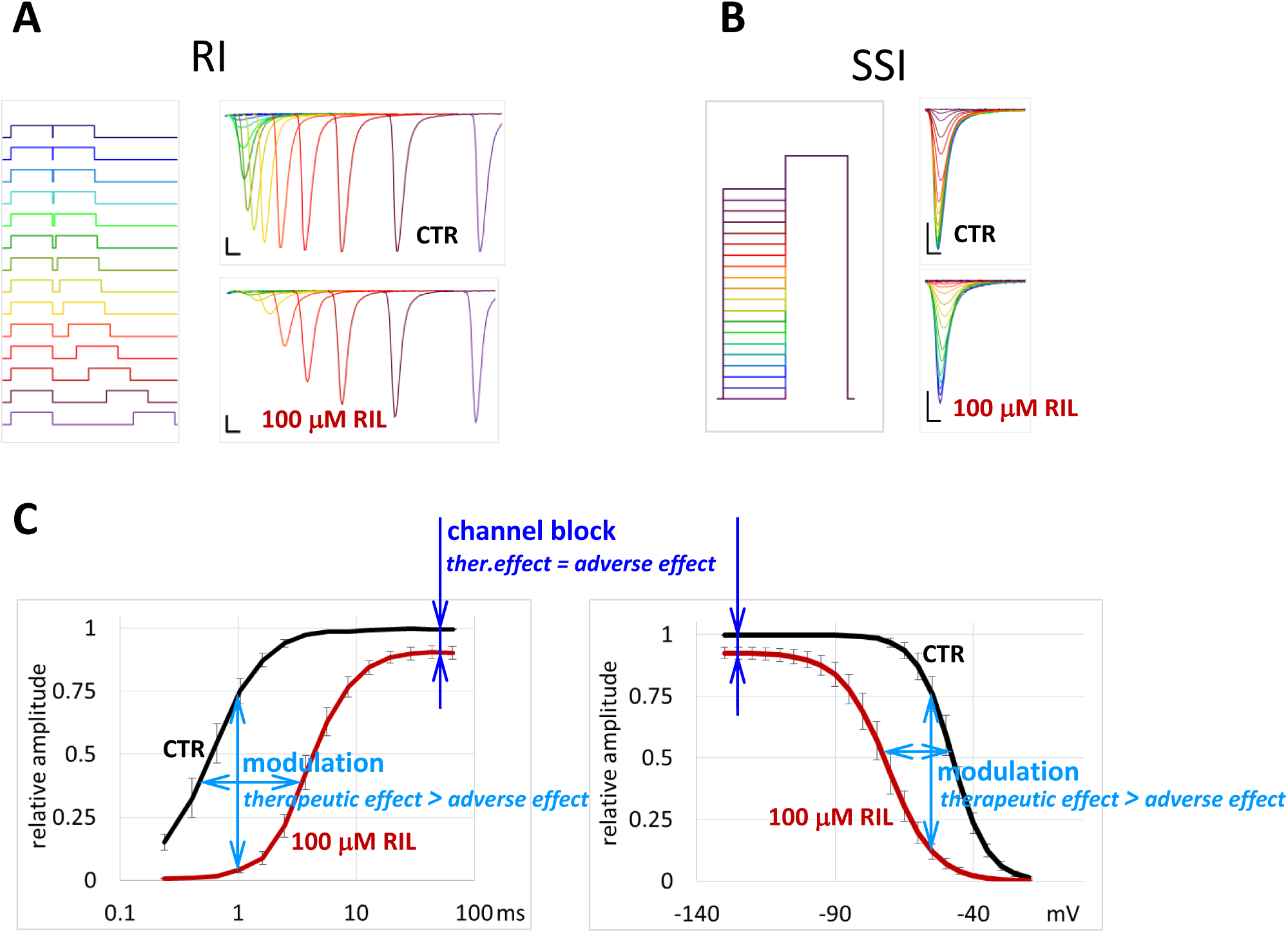
*The extent of channel block and modulation can be assessed using ***RFI*** and ***SSI*** protocols.* **A** Illustration of the first 22 ms of the **RFI** protocol. Left panel indicates the arrangement of 10 ms long depolarizing pulses (−130 to −10 mV), right panel illustrates currents evoked by the 2^nd^ pulse in a cell in control solution and in the presence of riluzole, on linear time scale. Scale bars: 1 ms and 1 nA. **B** Illustration of the **SSI** protocol. Left panel shows the voltage protocol (10 ms pre-pulses from -130 to -30 mV in 5 mV increments, followed by a 10 ms test pulse to -10 mV). Right panel shows examples for currents evoked by the test pulse in control solution and in the presence of riluzole. **C** Assessment of channel block and modulation using the **RFI** (plotted on a logarithmic time scale) and **SSI** protocols. Amplitudes were normalized to the maximum amplitude of control; mean amplitudes were obtained as described in text. Resting channel block is observed when sufficient time has been spent at hyperpolarized membrane potential. The effect of modulation is seen by the shift of curves.

From the therapeutic point of view modulation is more desirable than channel block, because while channel block equally affects healthy and diseased cells, modulation much more depends on the membrane potential and activity pattern of the cell, therefore, it may preferentially affect pathological cells, which either have an impaired maintenance of resting membrane potential, or have developed hyperexcitability (as a result of traumatic injury, inflammation, ischemia, etc.). The standard **SSI** and **RFI** protocols are able to assess both resting channel block (seen as the extent of inhibition after the cell has spent sufficient time at negative holding potentials) and modulation (seen as the extent of shift of **SSI** and **RFI** curves) at the same time (Fig. 1).

The obvious question is, whether channel block and modulation can be separated? Are there sodium channel inhibitor compounds that are predominantly blockers, and ones that are predominantly modulators? Could we specifically search for effective modulators which are at the same time weak blockers? These compounds would be expected to cause less adverse effects but more therapeutic effect.

### Unique properties of riluzole

One good example for a favorable modulation vs. block profile is the neuroprotective drug, riluzole (Fig. 2), which has been found to show uncommonly strong state-dependence: strong shift of the **SSI** curve with minimal inhibition of current amplitude at hyperpolarizing membrane potential (Fig. 1). Both in our comparative study of 35 drugs (Lenkey et al., 2010), and in a meta-analysis of literature data (Lenkey et al., 2011), riluzole was found to be one of the compounds with the highest state-dependence (Fig. 3). In this current study we intended to understand the reason for this property: Is it simply due to a higher difference between resting-state and inactivated-state affinities, or can it be due to the fact, that modulation is the principal mechanism, and channel block is only secondary? Could one find SCI drugs that are predominantly modulators? If yes, these would be more selective for pathological conditions, and therefore would show less adverse effects.

**Fig. 2.**
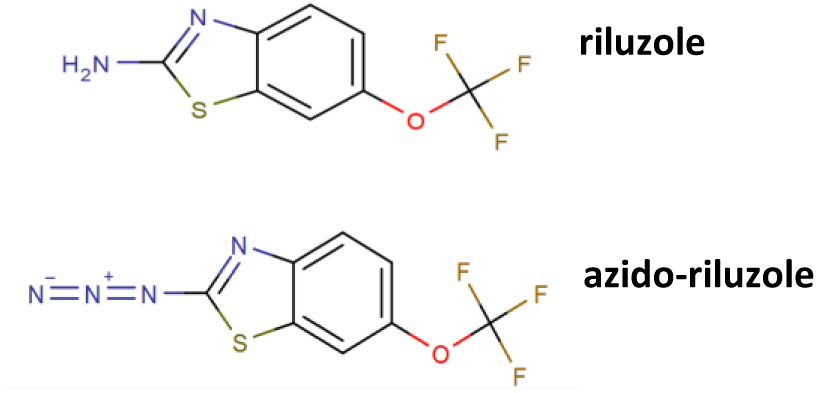
*Chemical structure of riluzole and azido-riluzole*.

The problem with separating modulation from block is that in experiments we can only distinguish conducting and non-conducting channels. For example a delayed recovery observed in a standard **RFI** protocol is usually thought to reflect dissociation of the drug, because drug-bound channels are generally considered non-conducting per se. However, if we suppose that drug binding does not necessarily exclude conduction, delayed recovery could also reflect the gating process itself, without dissociation (recovery itself must be delayed, according to the modulated receptor hypothesis). One way to test this is to eliminate one of the unknowns, the possibility of dynamic association and dissociation of the ligand during experiments. To accomplish this, we used a photoreactive riluzole analog, azido-riluzole (Fig. 2A), which can be induced to bind covalently to the channel by UV illumination.

**Fig. 3.**
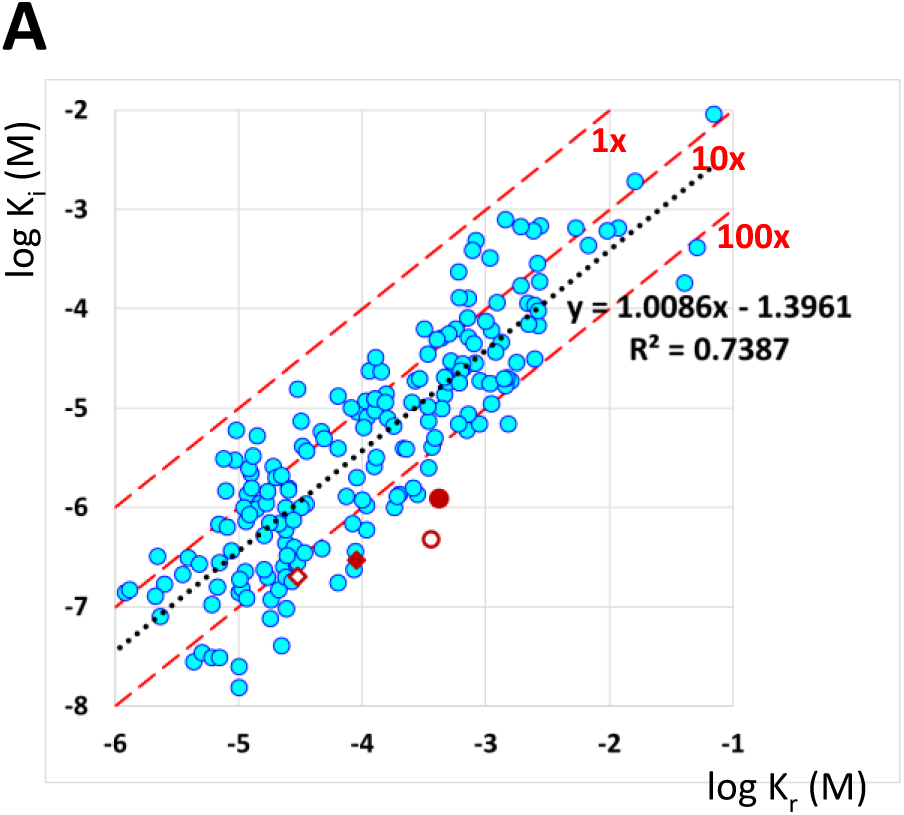
*State-dependence of riluzole in comparison with 122 SCI compounds.* Data are from Supplement #3 of (Lenkey et al., 2011); excluding Nav1.8-selective compounds. Resting and inactivated affinities are shown for 194 individual measurements of 122 compounds from 73 publications. Data were calculated from inhibition at different holding potentials, as described in (Bean et al., 1983); corrected and standardized, as described in (Lenkey et al., 2011). Data for riluzole are shown in red: closed diamond: (Benoit and Escande, 1991), open diamond: (Hebert et al., 1994), closed circle: (Lenkey et al., 2010), open circle: this study. The black dotted line is the linear regression line, with equation and correlation coefficient (R^2^). The average state-dependence, calculated from the y intercept of the linear regression line was 24.8-fold. Red dashed lines indicate K_r_ = K_i_; K_r_ = 10*K_i_; and K_r_ = 100*K_i_. The state-dependence of riluzole was above 100-fold in all studies.

### Photolabeling-coupled electrophysiology

The method of photoaffinity labeling (PAL) is conventionally used for target and binding site identification (Catterall, 2015; Dormán and Prestwich, 2000; Smith and Collins, 2015; Woll et al., 2016). However, the method can also be used to study direct effects of ligands (agonists, antagonists, inhibitors, modulators), when combined with electrophysiology. This allows real-time monitoring of the effect and its reversibility, both with and without photoactivation. Photolabeling-coupled electrophysiology has been used to study the interaction between binding site occupancy and gating, stoichiometry and cooperativity in different ligand-gated ion channels (Bhargava et al., 2012; Forman et al., 2007; Mortensen et al., 2014; Ruiz and Karpen, 1997; Zhong et al., 2008). In the case of sodium channels, our motivation was to test, whether the modulation of channel gating kinetics persisted after covalently binding and then washing out the photoreactive inhibitor compound. We reasoned that if modulation still persists after complete washout, then this state-dependent component of inhibition must come from a non-blocking modulation of sodium channels.

When designing a photoreactive sodium channel inhibitor, certain properties of the binding site must be taken into consideration. The binding site is located in the "inner vestibule", which is an aqueous cavity inside the pore region of the channel, surrounded by the S5 and S6 transmembrane segments of all four domains. It is limited by the selectivity filter from the extracellular side, and by the activation gate from the intracellular. Access to the inner vestibule is either through the activation gate during open state (hydrophilic pathway), or through the "fenestrations" directly from the membrane phase (hydrophobic pathway). Most drugs predominantly use the hydrophobic pathway, i.e., they can readily associate even without channel opening. Mutagenesis studies indicate that residues of S6 segments from domains I, III and IV contribute to the binding site (Mike and Lukacs, 2010), the most important being a Phe residue in DIV (F1764 in Nav1.2). Because of these properties of the binding site, there are some strict structural requirements to consider when designing a photoreactive SCI analog: Because the access pathway involves partitioning into the lipid bilayer, lipophilicity of the compound is crucial. There is a strong correlation between lipophilicity (logP) and affinity for SCI compounds, probably because SCI molecules accumulate within the membrane phase (Lenkey et al., 2010, 2011). In addition, the number of aromatic rings is strongly correlated with onset and offset rates and reversibility (Lazar et al., 2015; Lenkey et al., 2010). Finally, the size of the molecule is also crucial. One reason for the exceptional properties of riluzole might be its size; it has the lowest atom count (20) and van der Waals volume (164 Å^3^) among effective SCI compounds (Lazar et al., 2015; Lenkey et al., 2011). For these reasons, attaching one of the most commonly used groups, phenylazide, phenyldiazirine or benzophenone, would have been unadvisable. Fortunately, the aryl-amine group of riluzole can be changed into an aryl-azide, which is the smallest possible modification that produces a photoreactive analog. The azide group still alters important physicochemical properties of the compound, most importantly the charge distribution and lipophilicity, therefore the similarity of the effect of riluzole and azido-riluzole must be investigated. The compound was found to possess stability in the dark, and was highly reactive upon UV irradiation. Intermediates of photo-activated aryl-azides had sufficiently short lifetime (in the ps range) (McCulla et al., 2006), therefore binding should occur before the activated drug leaves the binding site by diffusion.

## Methods

### Cloning and stable cell line generation

To maximize recombinant sodium channel expression, the coding sequence of the rat NaV1.4 sodium channel was inserted into a modified pBluescript KS (Stratagene) vector (pCaggs IgG-Fc) capable to recombine with the murine Rosa26 BAC (Zboray et al., 2015). Briefly, NaV1.4 was inserted into the vector at *AscI* sites under the control of the Caggs promoter. Recombination to the *Rosa26* BAC was carried out by Recombineering (Muyrers et al., 1999; Zhang et al., 1998). Original BAC clone RP24-85I15 was derived from the BACPAC Resources Center (Children’s Hospital Oakland Research Institute). NaV1.4 BAC was transfected into CHO DUKX B11 (ATCC CRL-9096) suspension cells by Fugene HD (Promega) transfection reagent according to the manufacturer’s recommendations. Cell clones with stable vector DNA integration were selected by the addition of G418 antibiotic to the culture media (400 mg/ml) for 14 days.

### Cell culture and expression of recombinant sodium channels

CHO cells were maintained in Iscove's Modified Dulbecco’s Medium with 25mM HEPES and L-Glutamine (Lonza) supplemented with 10% v/v fetal calf serum, 200mM L-glutamine, 100 U/ml of penicillin/Streptomycin, 0.5 mg/mL Geneticin(Life Technologies) and 2% ProHT Supplement (Lonza). Cells were plated onto 35 mm petri dishes and cultured for 24-36 hours. Prior to experiments cells were dissociated from the dish with trypsin-EDTA, centrifuged and suspended into the extracellular solution.

### Materials

All chemicals were obtained from Sigma-Aldrich. Azido-riluzole was synthesized by SONEAS Research Ltd. Budapest, Hungary

### UV photoactivation

The recording chamber of a Port-a-Patch system (Nanion, Munich, Germany) was customized to accommodate a 400 μm diameter quartz optic fiber, which was placed 3–4 mm above the recorded cell. The original perfusion manifold was replaced by a custom manifold positioned to the side of the recording chamber opposite to the waste removal. Solution exchange was complete within 1-2 s. UV light was applied for 180 s, using a 310 nm fiber coupled LED (Mightex FCS-0310-000), with 40 μW intensity.

### Electrophysiology

Whole-cell currents were recorded from cells voltage clamped at −130 mV using an EPC10 plus amplifier, and the PatchMaster software (HEKA Electronic, Germany). During cell catching, sealing, and whole-cell formation, the PatchControl software (Nanion, Munich, Germany) commanded the amplifier and the pressure control unit. Currents were filtered at 10 kHz, and digitized at 20 kHz. The intracellular solution contained (mM): 50 CsCl, 10 NaCl, 60 CsF, 20 EGTA, 10 HEPES; pH 7.2 (adjusted with 1M CsOH). The resistance of borosilicate chips was 2.0 – 3.5 MΩ. Cells were continuously perfused with extracellular solution containing (mM): 140 NaCl, 4 KCl, 1 MgCl2, 2 CaCl2, 5 D-Glucose and 10 HEPES; pH 7.4 (adjusted with 1M NaOH).

### Data analysis

Curve fitting was done in Microsoft Excel, using the Solver Add-in. Steady-state inactivation (**SSI**) curves were fitted using the Boltzmann function: *I* = *I_max_*/{1+exp[(*V_p_*-*V_1/2_*)/-*k*]), where V_p_ is the pre-pulse potential, V_1/2_ is the voltage where the curve reaches its midpoint, and k is the slope factor. Recovery from inactivation (**RFI**) data were fitted by exponential function. We noticed that a simple exponential function did not adequately fit data points, the fit much improved when the exponential equation was on the second power, and further improved on the third power: *I* = *A*\*[1-exp(-*t_ip_*/ *τ*)]^3^, where A is the amplitude, and *t_ip_* is the duration of the interpulse interval. The time constant of best fit equations changes with the power, for example the same control recovery curve was fitted with *τ* = 1.09 ms (1^st^ power), *τ* = 0.62 ms (2^nd^ power), and τ = 0.48 ms (3^rd^ power). In order to keep time constants comparable we fixed the exponent at the value of 3 for all curves during fitting. In the presence of azido-riluzole adequate fitting required a second exponential component, therefore these data were fit with the following equation: *I* = *A_1_*\*[1-exp(-*t_ip_*/ *τ_1_*)]^3^ + *A_2_*\*[1-exp(-*t_ip_*/ *τ_2_*)].

Averaging was not done by averaging data points across individual cells, because that would have resulted in an erroneously decreased slope of averaged curves. Instead, for each individual cell data points were fitted separately, and curves were reconstructed from the averaged parameters of equations (time constants and amplitudes in the case of **RFI** curves, *V_1/2_* values and slope factors in the case of Boltzmann equations to reconstruct **SSI** curves). Error bars show SEM for original data points. All data are presented as mean ± SEM for the indicated number of experiments (n). Significance levels were calculated using paired Student's *t* test. Chemical property prediction and calculation was done using JChem for Excel 15.4 software from ChemAxon (http://www.chemaxon.com).

## Results

### Inhibition of NaV1.4 sodium channels by riluzole

The effect of riluzole on the gating equilibrium kinetics was studied using the steady-state inactivation (SSI) and the recovery from inactivation (**RFI**) protocols. In the **SSI** protocol the membrane potential was set for 10 ms to pre-pulse potentials between −130 mV and −20 mV in 5 mV increments, which was followed by a 10 ms test pulse to -10 mV, in which the availability of the channel population was assessed (Fig. 1). Data points were fit with the Boltzmann function, from which the half inactivation voltages (V_1/2_) values were determined. In control experiments the V_1/2_ was 56.3 ± 3.76 mV, which was shifted to the hyperpolarized direction by -27.5 ± 3.95 mV by 100 mM riluzole (Fig. 4A). The V_1/2_ value did not always recover to its original value upon washout, because – as it is well documented in the literature – it tends to undergo a spontaneous left shift during whole-cell recording. For this reason, the shift caused by drugs may be somewhat overestimated. This drawback, however was absent in the **RFI** protocol, where the time constant of recovery always faithfully regained its control value upon washout. The **RFI** protocol consisted of two equal 10 ms depolarizations to -10 mV, separated by a hyperpolarizing gap (-130 mV) with logarithmically increasing duration (multiplied by 1.5) between 0.1 ms and 65.7 ms (Fig. 1). In the presence of 100 μM riluzole the time constant was increased 6.33-fold (range: 5.27- to 9.65-fold): while drug-free channels recovered from inactivated state with the time constant of 0.49 ± 0.09 ms, in the presence of riluzole the time constant was 2.97 ± 0.50 ms (p = 4*10^-7^, n = 7) (time constants are for the exponential on the third power, see Methods). At the same time, the decrease in amplitude was minimal, it decreased to 87.5 ± 4.78% of the original amplitude (p = 0.03). Individual measurements and the average of fitted curves are shown in Fig. 4.

**Fig. 4.**
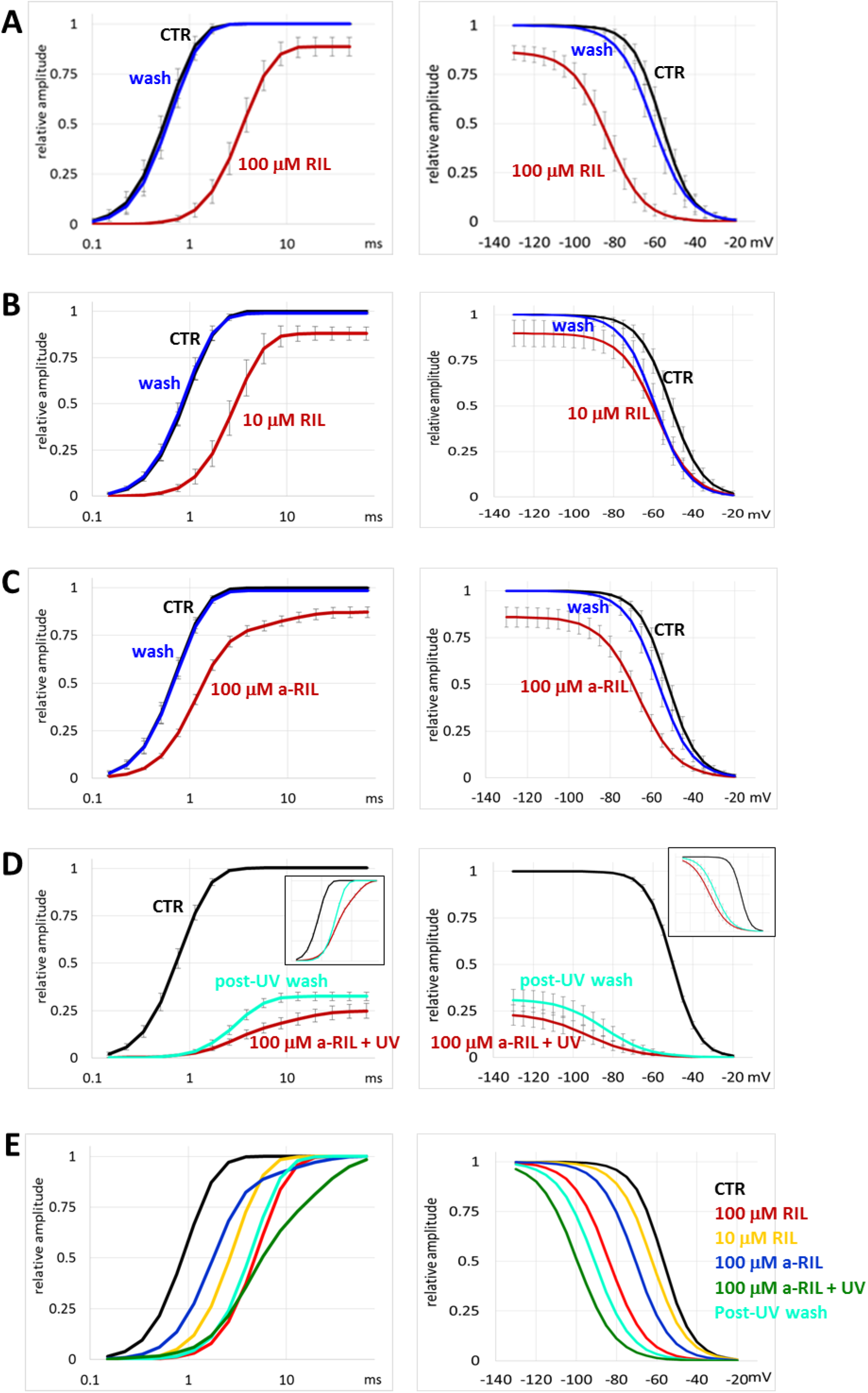
*The effect of μM riluzole and μM azido-riluzole in the **RFI** and **SSI** protocols.* Black, red and blue lines indicate control, drug and washout, respectively**. A** 100μM riluzole. **B** 10 μM riluzole. **C** 100 μM azido-riluzole. **D** 100 μM azido-riluzole during UV-irradiation, and after washout. Turquoise lines indicate washout after azido-riluzole perfusion and UV-irradiation. Insets show curves normalized to their own maxima. Averaging was done as described in Methods. **E** Comparison of the effects of all five treatments. All curves were normalized to their own maxima.

In order to monitor the onset and offset of the effect, we used a three-pulse train (**3PT**) protocol (Fig. 5), which was designed to indicate changes both in the gating kinetics and gating equilibrium. The first depolarizing pulse measured resting inhibition, because it was evoked from a prolonged hyperpolarized membrane potential (263 ms at −130 mV). The second was designed to indicate changes in the recovery kinetics, and it followed the first one after a 5 ms hyperpolarizing gap (−130 mV). This duration enables drug-free channels to recover completely from inactivation (99 ± 0.004%), while in the presence of 100 μM riluzole the recovery was incomplete (57.7 ± 8.7%). A drug that modulates recovery kinetics, should show a selective inhibition of the 2^nd^ pulse as compared to the 1^st^ one. Finally, the 3^rd^ pulse was designed to indicate changes in gating equilibrium. Before this 3^rd^ pulse the holding potential was elevated to the approximate V_1/2_ value (set individually for each cell, ranging between −75 and −50 mV) for 50 ms. The amplitude of this 3^rd^ pulse was a very sensitive indicator of changes in the voltage-dependence of inactivation. Three-pulse trains were evoked every 333 ms. Using this protocol, we could measure the three important properties (resting inhibition, modulation of gating kinetics, modulation of gating equilibrium) of the effect of SCIs in parallel and with reasonable time resolution. Higher frequency would not improve time resolution, because it was limited by the rate of solution exchange (~1 – 2 s). The time course of the onset and offset of inhibition is shown in Fig. 5 for the three pulses in parallel. The ratios of 2^nd^ / 1^st^ and 3^rd^ / 1^st^ pulse-evoked amplitudes are even more informative regarding the extent of modulation; they indicate the drug's potency for affecting gating kinetics and gating equilibrium, respectively. We show these ratios, calculated from the average traces for 100 μM and 10 μM riluzole, as well as for azido-riluzole with and without UV irradiation in Fig. 5E. Note that riluzole was much more effective as a modulator than as a resting state blocker.

**Fig. 5.**
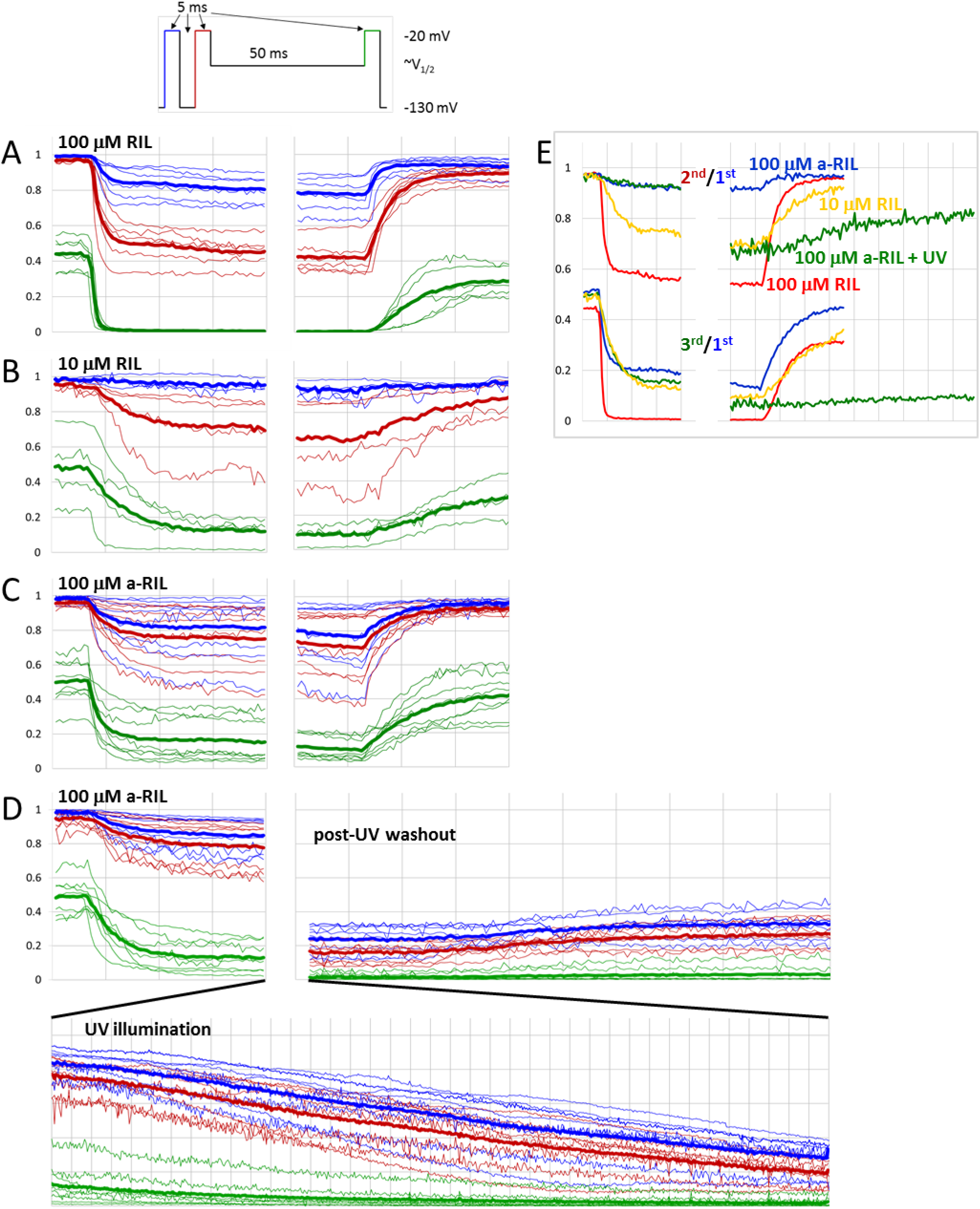
*Onset and offset of drug effects in the 3PT protocol*. Plots of peak amplitudes for the 1^st^, 2^nd^, and 3^rd^ pulse-evoked currents are shown in blue, red and green lines, respectively. Thin lines show measurements from individual cells, thick lines show averages. Recording from individual cells were aligned to the moment of wash-in and wash-out for comparability, with 0 to 767 s of recording not shown between the onset and offset. Ten control trains (3.3 s) are shown before drug applications, and the last 20 trains (6.6 s) of drug application before wash-out. **A** 100 μM riluzole, **B** 10 μM riluzole, **C** 100 μM azido-riluzole, **D** 100μM azido-riluzole with UV irradiation (shown on a contracted time scale below onset and offset). **E** Plot of 2^nd^ / 1^st^ pulse-evoked (upper curves) and 3^rd^ / 1^st^ pulse-evoked (lower curves) amplitude ratios for the onset and offset of: 100 μM riluzole (red lines), 10 μM riluzole (yellow lines), 100 μM azido-riluzole (blue lines), and 100μM azido-riluzole with ~3 min UV irradiation between onset and offset. Vertical gridlines indicate 5s in all figures.

This was clearly observable even at 10 μM concentration, where the onset rate was lower, and the inhibition somewhat smaller, but the considerable modulation (see 2^nd^ / 1^st^ and 3^rd^ / 1^st^ pulse-evoked amplitude ratios in Fig. 5B and E) as compared to the minimal resting block (see the minimal effect on the 1^st^ pulse-evoked current) shows the peculiar nature of inhibition by riluzole.

Application of UV irradiation for 180 s did not change the inhibition by riluzole in any of the three protocols. Similarly, UV irradiation did not alter control values (data not shown).

### Inhibition by azido-riluzole

Without UV irradiation 100 μM azido-riluzole caused a weaker shift in both **SSI** and **RFI** curves than riluzole, although the resting state inhibition was similar (Fig.4B). This indicates that it is an equipotent blocker, but a weaker modulator than riluzole. In addition, the onset and offset of inhibition were slower, as seen in the **3PT** protocol (Fig. 5C). This was expected, because access to the binding site should be faster in the case of the more lipophilic, and less polar riluzole, which also has a more uniform charge distribution. It is likely that the concentration that determines the occupancy of SCI binding sites is not the aqueous concentration of the drug, but rather the concentration within the membrane phase, which must be lower for azido-riluzole than for riluzole. In terms of modulation of the gating equilibrium, 100 μM azido-riluzole was roughly equi-effective with 10 μM riluzole, as seen by the shift of the **SSI** curve (Fig.4B and C), and the 3^rd^ / 1^st^ pulse evoked current amplitude ratio (Fig.5E). The V_1/2_ was shifted to the hyperpolarized direction by −14.0 ± 2.21 mV (Fig. 6). In terms of modulation of gating kinetics, azido-riluzole had even weaker effect than 10 μM riluzole, as seen in the **3PT** protocol, where there was hardly any difference between the inhibition of the 1^st^ and 2^nd^ pulse-evoked currents. (Fig. 5C and E). In addition, **RFI** curves were only moderately shifted; the fast time constant of recovery increased only 1.6-fold (range: 1.33- to 2.36-fold; p = 1*10^-4^, n = 7) (Fig.4B and C). In addition to the delayed fast component, a second, slow component appeared, the time constant of which was 7.37 ± 1.23 ms (range: 4.65 to 14.2 ms), and its contribution to the amplitude was 17.6 ± 3.8% (range: 11.1 to 35.2%) (Fig. 6B).

### The effect of UV-illumination in the presence of azido-riluzole

We applied a single 180 s UV irradiation in the presence of 100 μM azido-riluzole while running the **3PT** protocol, which resulted in a slow continuous decrease of 1^st^ pulse evoked sodium current amplitudes, to 25.3 ± 3.6% of its initial value (Fig. 5D). The slow decay depended on the presence of UV light: when illumination was suspended, the decay was also temporarily halted. The inhibition recovered to 32.5 ± 3.8% of the initial value upon washout, the roughly 7% difference probably reflects dissociation and washout of unactivated azido-riluzole. By the end of azido-riluzole treatment, UV irradiation and washout, thus 67.5% of the control current amplitude was irreversibly inhibited by the covalently bound azido-riluzole. The inhibition persisted throughout the rest of the experiment, for up to 20 minutes post-UV exposure. Currents evoked by 2^nd^ pulses were irreversibly reduced to 25.5 ± 2.8%, while currents evoked by the 3^rd^ pulse to 2.78 ± 1.62% of the pre-UV amplitude.

One crucial question was, whether covalently bound azido-riluzole causes only channel block, or also modulation. In this situation when unbound azido-riluzole had already been washed out, delayed recovery cannot reflect the process of dissociation, only modulated gating. Similarly, a shifted **SSI** curve cannot be caused by state-dependent affinity, because no association and dissociation can occur, therefore the only explanation must be modulation. If one can see signs of modulation by covalently bound inhibitor, it proves that non-blocking modulation is possible, and therefore, it might be responsible for a significant fraction of inhibition in the case of non-covalently bound riluzole as well. This could explain the peculiar behavior of riluzole (small resting state inhibition with strong modulation).

As it can be seen in Fig. 4D, the remaining component of the sodium current was strongly modulated by the covalently bound inhibitor (there was no perfused inhibitor present). The extent of modulation was close to the values observed in the presence of 100 μM riluzole. The **SSI** curve was shifted by −42.8 ± 4.33 mV post-UV in the presence of 100 μM azido-riluzole, and was still shifted by-34.3 ± 6.93 mV after washout. Recovery kinetics was clearly bi-exponential in the presence of azido-riluzole, the fast component increased 3.99-fold (range: 2.46- to 6.14-fold; p = 7.8*10^-4^, n = 8), and a slow component emerged with a time constant of 12.6 ± 1.31 ms (range: 7.2 to 17.7 ms), contributing 52.1 ± 3.18% to the amplitude (range: 38.7 to 62.4%). After washing out unactivated azido-riluzole, the time course of recovery again could be reasonably well fitted with a single exponential component (on the third power), which probably indicates that the slow component was due to azido-riluzole dissociation. The time constant after washout was 4.48-fold higher than the time constant of control curves (range: 2.44 to 7.57-fold change; p = 1.7*10^-5^, n = 8).

Note that – just as in the case of 100 μM riluzole, see Fig. 4A and D – inhibition was close to 100% at gap durations < 1 ms. This indicates close to full occupancy of binding sites, nevertheless, 32.5 ± 3.8% of the control current was able to flow through channels with covalently bound inhibitors. Binding to areas of the channel other than the primary binding site (known as the "local anesthetic receptor") might have occurred, however, this could not account for modulation. Mutation of the key residue of the primary binding site, Phe1579 abolished modulation ( Szabo et al., bioRxhiv), similarly to what have been observed in other laboratories, where this mutation selectively abolished use-dependent inhibition (Ragsdale et al., 1994), the shift of SSI curves (Hanck et al., 2009) as well as the effect of SCIs on voltage sensor movement (Hanck et al., 2009; Muroi and Chanda, 2009). Similarly, in the **SSI** curves inhibition was close to 100% at membrane potentials >-50 mV. Full inhibition can only be caused by full occupancy of binding sites, which implies that at hyperpolarized potentials conduction occurred in spite of the presence of a covalently bound inhibitor at the binding site.

## Discussion

We have performed a photolabeling-coupled patch-clamp study on hNav1.4 sodium channels, and the effect of a novel photoactive inhibitor, azido-riluzole. Our method allowed not only to monitor the extent of inhibition during the process of photolabeling, but also to assess the contribution of channel block and channel modulation.

Riluzole is unique among SCIs in its high ability to modulate and relative low potency to cause channel block. We hypothesize that this may be due to the small volume, high lipophilicity and neutrality of this molecule, which may make compounds with such physicochemical properties promising therapeutic drugs. Most importantly we wanted to test if channel block (which unavoidably leads to adverse effects) can be separated from modulation of channel gating (which has the potential to cause selective inhibition of pathological activity); *i.e.*, if modulation can exist without channel block.

In order to answer this question we needed to eliminate the unknown of association/dissociation dynamics of the ligand from the experiments. Interpretation of kinetic experiments is complicated by the simultaneous and mutually interacting processes of channel gating and drug binding/unbinding. If one can make experiments with covalently bound inhibitor molecules, this ambiguity can be excluded.

We performed experiments with the photoactive analog of riluzole, azido-riluzole. The compound could be activated by UV irradiation at 310 nm wavelength, which – at least for the 180 s duration we used in our experiments did not cause noticeable damage to the cells. This indicates that azido-riluzole could be used in in vivo experiments as well. We demonstrated that UV illumination increased its potency (as if it was applied at a higher concentration), and made it to be irreversible on the time scale of an average electrophysiology experiment. Upon UV irradiation a virtual increase in affinity occurs, which allows this compound to be used in experiments where localized effects of sodium channel inhibition are to be studied.

The compound azido-riluzole did not qualify as the exact replica of riluzole, with the only difference of being photoactive. The necessary changes in chemical composition caused it to have less affinity, furthermore it became more of a blocker and less of a modulator as compared to riluzole. Nevertheless, it still maintained a definite modulatory component of inhibition. The principal question was, if it would still maintain its ability to modulate after having bound covalently to its binding site within the channel.

Significantly, after photoactivation, covalent binding to the channel, and subsequent wash-out of the unbound fraction, the bound fraction of azido-riluzole was still able to exert a modulatory effect that resembled the modulation caused by 100 μM riluzole. We could conclude that non-blocking modulatory effect on sodium channels is possible, and it is the most likely explanation for the extraordinary properties of inhibition caused by the parent molecule, riluzole as well. We hypothesize that this special kind of inhibition is due to the special physicochemical properties of riluzole, namely its small volume, neutrality and lipophilicity. This hypothesis opens the way to the search for other non-blocking modulatory SCI drugs, which are expected to have stronger therapeutic and less severe adverse effects.

## Acknowledgements

This work was supported by the Hungarian Brain Research Program (KTIA-NAP-13-2-2014-002), and The National Research, Development and Innovation Office (VKSZ_14-1-2015-0052).

